# Telomere-to-telomere human DNA replication timing profiles

**DOI:** 10.1101/2022.03.28.486072

**Authors:** Dashiell J. Massey, Amnon Koren

## Abstract

The spatiotemporal organization of DNA replication produces a highly robust and reproducible replication timing profile. Sequencing-based methods for assaying replication timing genome-wide have become commonplace, but regions of high repeat content in the human genome have remained refractory to analysis. Here, we report the first telomere-to-telomere replication timing profiles in human, using the T2T-CHM13 genome assembly and sequencing data for five cell lines. We find that replication timing can be successfully assayed in centromeres and large blocks of heterochromatin. Centromeric regions replicate in mid-to-late S-phase and contain replication-timing peaks at a similar density to other genomic regions, while distinct families of heterochromatic satellite DNA differ in their bias for replicating in late S-phase. The high degree of consistency in centromeric replication timing across chromosomes within each cell line prompts further investigation into the mechanisms dictating that some cell lines replicate their centromeres earlier than others, and what the consequences of this variation are.

## Introduction

Eukaryotic DNA replication initiation is organized in space and time, reflecting a reproducible DNA replication-timing program^1^. In general, late replication appears to be associated with a more repressive chromatin state: late-replicating regions tend to localize to the nuclear periphery^2,3^ and to broadly associate with the condensed “B” compartment in chromatin conformation capture assays^4,5^. Likewise, genes in late-replicating regions often have lower expression^6,7^, with corresponding histone methylation^8,9^ and deacetylation^8,10^, than genes in early-replicating regions. Constitutive heterochromatin, which is gene-poor and highly-condensed, is often described to be late replicating^11-13^, although direct visualization of nascent DNA by microscopy indicates that there are five distinct waves of replication initiation during S phase, with euchromatic replication primarily occurring during the first wave^2^. This suggests that heterochromatin replication timing is likely more complicated than currently appreciated, and potentially points to the existence of distinct heterochromatin subtypes that differ in their replication timing.

Existing methods for measuring replication timing at genome scale^14^ are sequencing-based, making them reliant on the quality of reference genome assemblies. Notably, the current human reference genome (GRCh38/hg38) contains 151Mb of unresolved gaps, represented as multi-megabase arrays of unknown sequence^15^. Thus, these regions – which include large pericentromeric regions on chromosomes 1, 9, and 16 and the entire p-arms of the five acrocentric chromosomes (chr13, chr14, chr15, chr21, chr22) – have been refractory to whole-genome analyses, including those of replication timing. In addition, hg38 contains statistically modeled sequences for the centromeric α-satellite DNA, which were designed as decoys for sequence alignment rather than to reflect the true linear sequence of these arrays^16^.

We previously reported^17^ that these centromeric sequence models in hg38 enabled preliminary analysis of replication timing for the majority of human centromeres. We found consistent evidence of replication-timing peaks within centromeric regions, suggesting that centromeres contain replication origins. We further demonstrated that centromeric replication occurs during mid-to-late S-phase and that its timing is highly divergent among cell lines. However, because the decoy sequences in hg38 were not linear assemblies of the centromeres, we were unable to analyze the precise locations of these peaks.

Here, we report telomere-to-telomere replication timing profiles across all autosomes and the X chromosome. Using the telomere-to-telomere human genome assembly T2T-CHM13, recently published by the Telomere-to-Telomere Consortium^15^, we provide the first report of replication timing of constitutive heterochromatin in the context of the whole genome. The linear sequences for the centromeres in this genome assembly further enabled us to revisit and reaffirm our previously conclusions based on hg38, while also analyzing the locations of centromeric replication initiation sites.

## Results and Discussion

### Telomere-to-telomere replication timing profiles

In our prior analysis^17^, we generated replication timing profiles for five cell lines – the apparently healthy lymphoblastoid cell line GM12878, the embryonic kidney cell line HEK293T, the ovarian carcinoma cell line A2780, and the breast cancer cell lines HCC1143 and HCC1954 – by whole-genome sequencing of G_1_- and S-phase populations isolated by fluorescence-activated cell sorting (FACS). The G_1_-phase fraction was used to define variable-size uniform-coverage genomic windows, accounting for sequencing biases and copy-number variants, and then sequencing read depth was assessed for the S-phase fraction. After S/G_1_ normalization, fluctuations in S-phase read depth reflect only the effects of replication timing, such that early-replicating regions are more highly represented relative to late-replicating regions^18^.

T2T-CHM13 is a gapless human genome assembly for CHM13-hTERT, a telomerase reverse transcriptase-transformed cell line derived from a complete hydatidiform mole with a stable 46, XX karyotype^15^. Hydatidiform moles are formed during fertilization and contain only DNA from the sperm; thus CHM13-hTERT is homozygous, reducing the complexity of genome assembly. T2T-CHM13 was assembled from long-read PacBio circular consensus sequencing and polished by with a combination of other short- and long-read sequencing methods. To assess whether this new assembly could be used to study the replication timing of heterochromatin, we generated replication timing profiles from the same sequencing libraries, re-aligning the sequencing reads for each cell line to T2T-CHM13. The resulting replication timing profiles were nearly gapless, with only the rDNA loci remaining as unresolved (Figure 1). (We note that CHM13-hTERT has an XX karyotype, as do all five cell lines studied. Thus, we did not consider the Y chromosome.) We validated these replication-timing profiles by comparison to the hg38-based replication timing profiles, using the UCSC Genome Browser liftOver tool to convert between hg38 and T2T-CHM13 coordinates. The profile for each cell line was virtually identical (r > 0.999) between genome builds for regions that could by successfully “lifted over”. Notably, this approach for inferring the replication timing of heterochromatic regions necessitated the analysis of a G_1_ control sample and was not amenable to FACS-free inference of replication timing from genome sequence data^19^ (Supplementary Fig. 1).

Our telomere-to-telomere profiles revealed the replication timing of several large regions previously excluded from genomic analysis. This included the entire p-arms of the acrocentric chromosomes (except for the rDNA loci) and the large pericentromeric satellite arrays on chromosomes 1, 9, and 16. The replication timing profiles in each of these regions showed similar structure to the profiles for other genomic regions, with distinct local maxima and minima of varying amplitudes (Figure 2a, b; Supplementary Fig. 2). Annotation of these new sequences^20^ indicated that these regions include several multi-megabase repeat arrays of distinct satellite sequences, including human satellite 1 (HSat1; 4.9Mb on chr13p), human satellite 2 (HSat2; 13.2Mb on chr1q, 12.7Mb on chr16q), human satellite 3 (HSat3; 27.6Mb on chr9, 8Mb on chr15p), and β-satellite (1.9Mb on chr22p). Within these larger satellite arrays, HSat1 appeared to replicate in mid-S phase, while HSat2 and HSat3 were later-replicating; we further characterize the replication timing of each satellite family, across all family members genome-wide, below.

**Figure 1.**
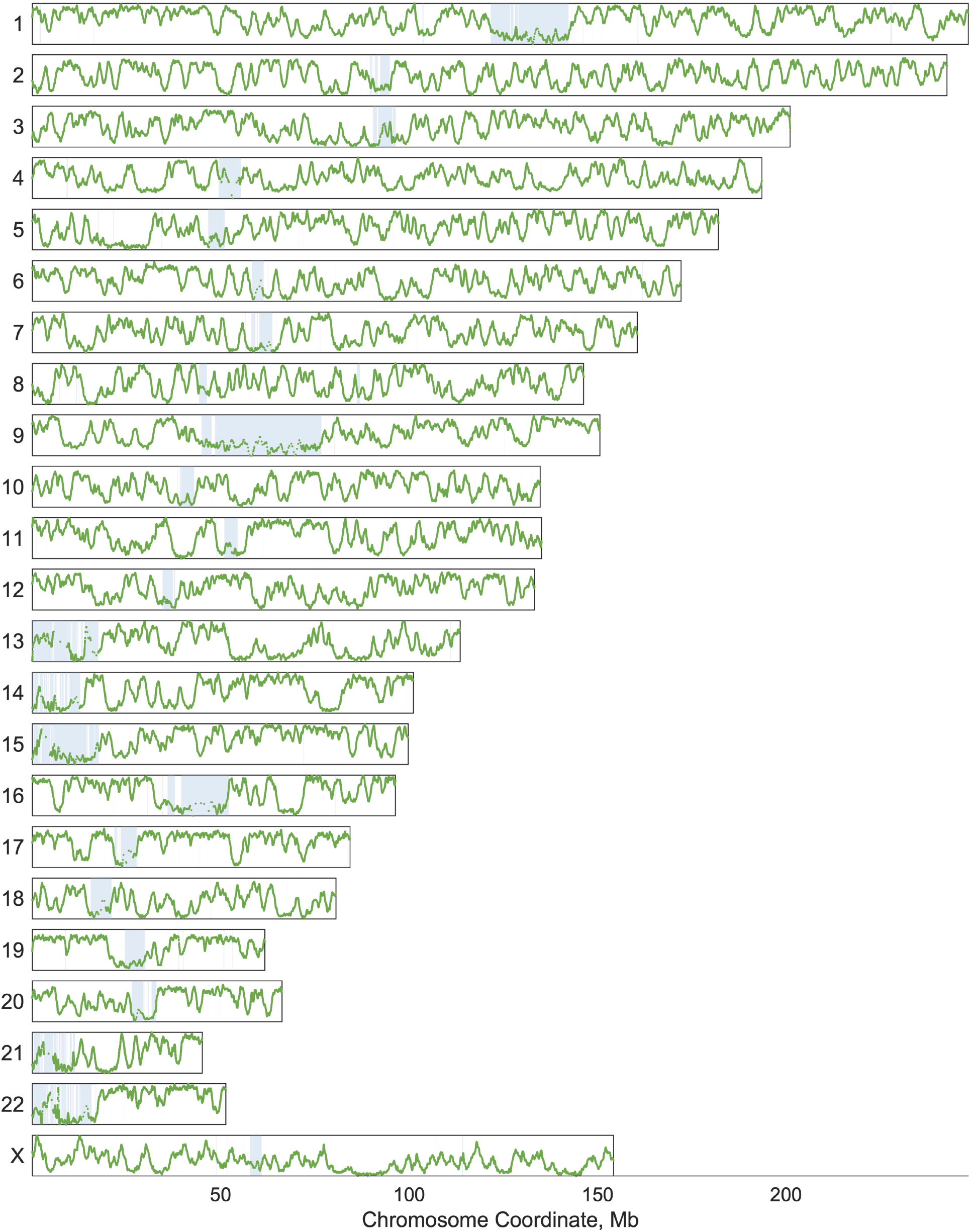
Telomere-to-telomere replication timing profiles for all autosomes and chromosome X. Regions larger than 5Kb that are new in T2T-CHM13 are indicated with the blue boxes. The replication-timing profile for GM12878 is shown.

**Figure 2.**
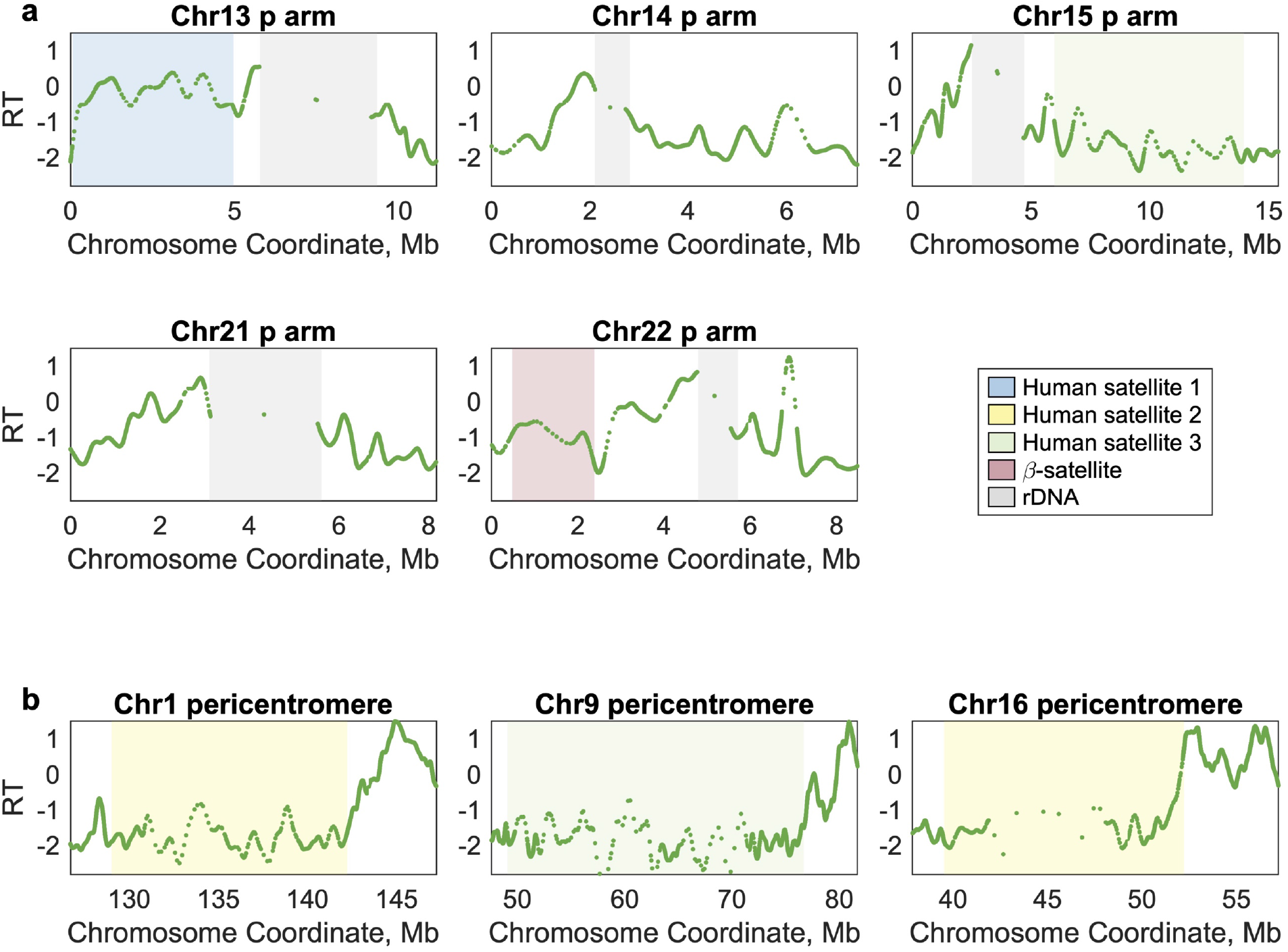
Replication timing (RT) of previously unresolved regions of the human genome. **a** RT profiles for the six acrocentric p-arms. rDNA arrays (gray) remain as gaps in the profile. **b** RT profiles for the large heterochromatin arrays neighboring the centromeres on the q-arms of chromosomes 1, 9, and 16. The RT profile for the lymphoblastoid cell line GM12878 is shown for each region.

Next, we visualized the centromeric regions. Using hg38, we previously reported that each centromeric region contains multiple replication timing peaks and that centromeric replication timing peaks were not particularly late relative to the rest of the genome^17^. Although the linear centromeric sequences in T2T-CHM13 completely replace the decoy sequences in hg38, these results were reproduced here (Figure 3; Supplementary Fig. 3). Additionally, we were able to meaningfully identify the locations of these local maxima within centromeric regions and to analyze their timing, as we present below. Furthermore, satellite repeat elements within T2T-CHM13 centromeric regions are well-annotated^20^, enabling us to characterize the replication timing of the rapidly-evolving centromere-specific α-satellite DNA, which is present as canonical higher-order repeat arrays (HORs), divergent higher-order repeat arrays, and α-satellite monomers (presented in Figure 4). Although many of the centromeric regions contain multiple HORs, only a subset is observed to bind kinetochore proteins and function in active centromere assembly^21^.

**Figure 3.**
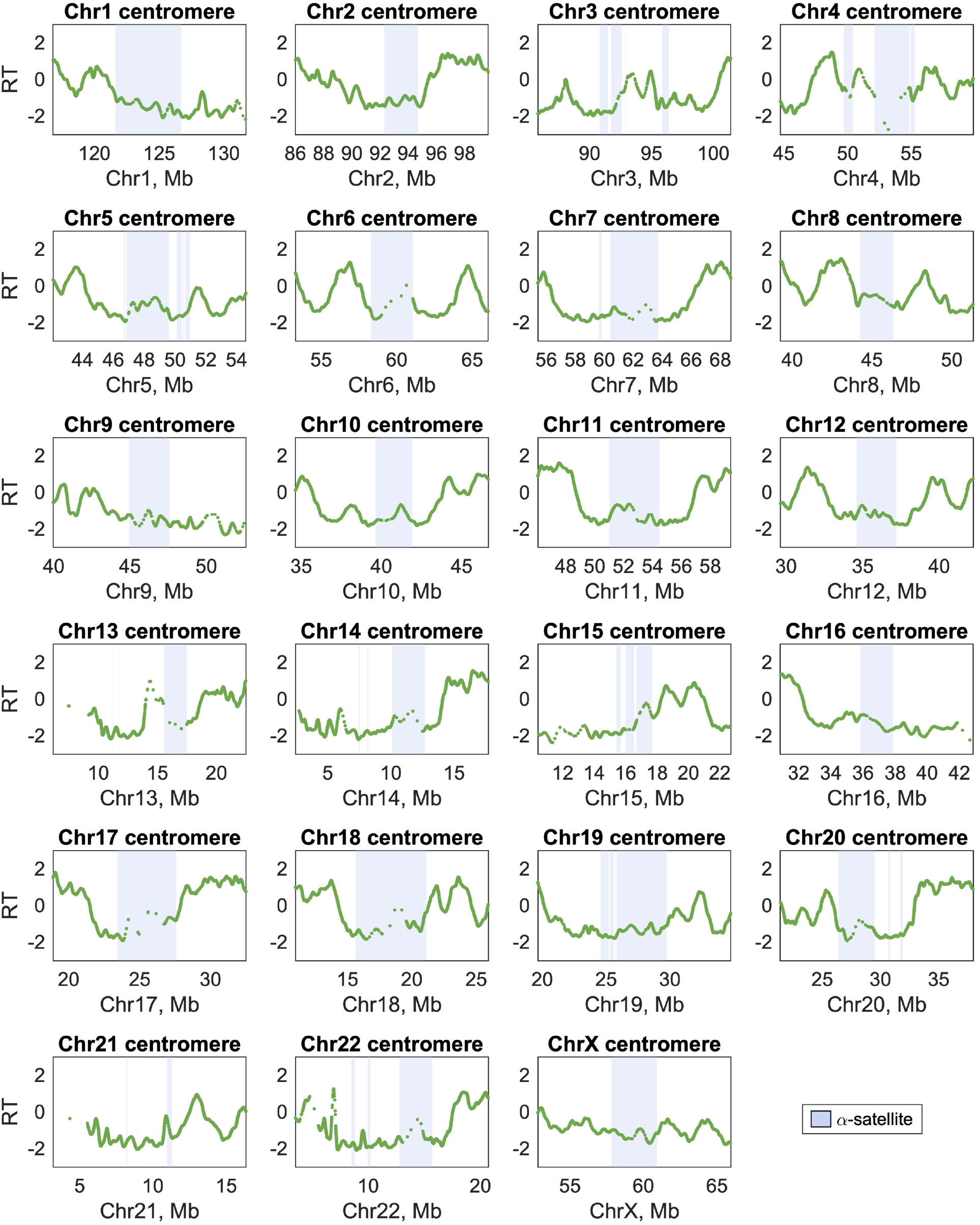
Centromeric replication timing (RT) of all human autosomes and chromosome X. The locations of α-satellite higher-order repeats on each chromosome, which scaffold active centromere assembly, are indicated in blue. For each chromosome, the entire region shown is annotated as centromeric. The RT profile for the lymphoblastoid cell line GM12878 is shown for each region.

**Figure 4.**
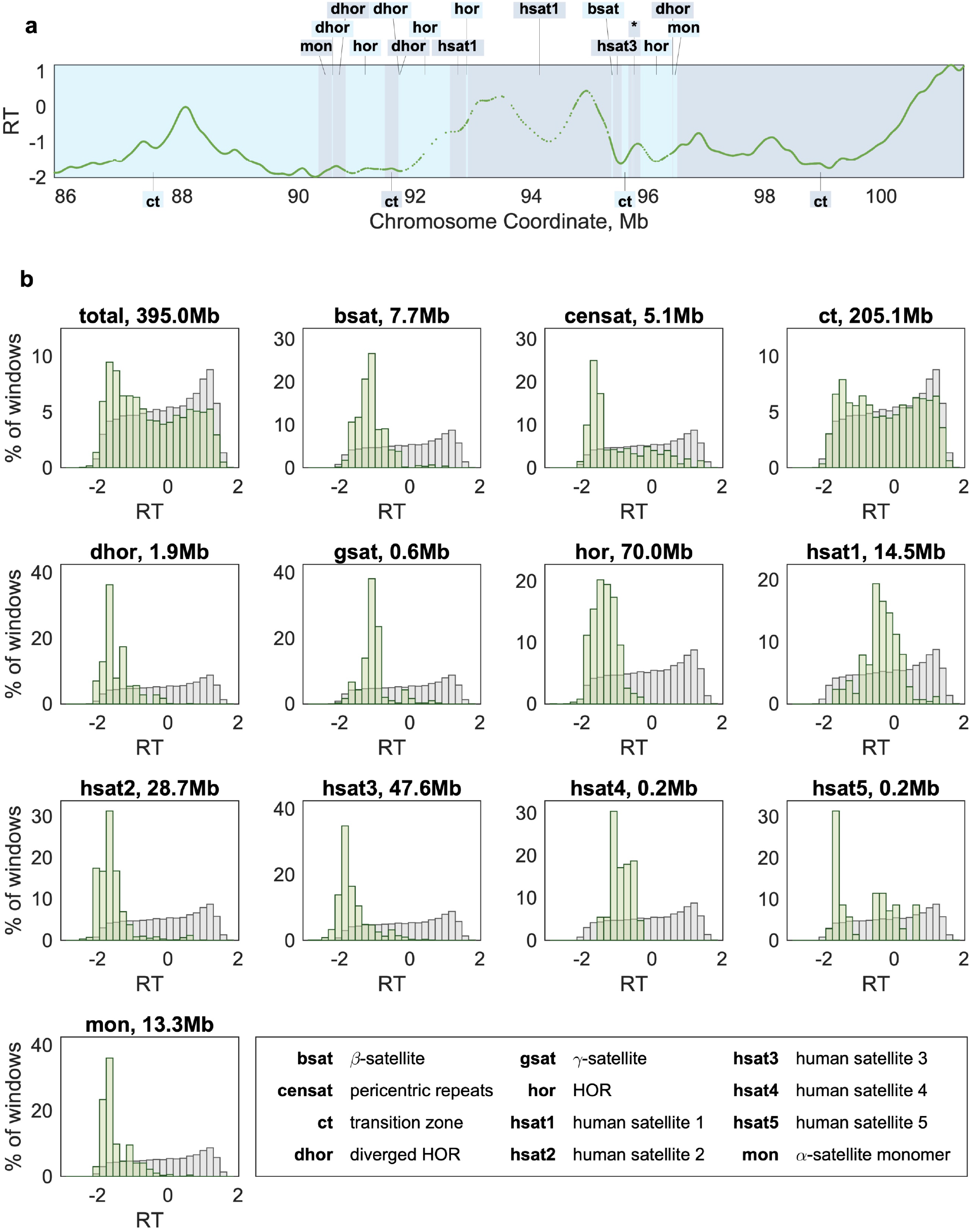
Replication timing (RT) bias of different satellite sequence elements. **a** The centromeric region of chromosome 3 is shown. Neighboring sequence elements are denoted in alternating colors. The 200kb region indicated with an asterisk contains 11 sequence elements. **b** For each sequence element category, the distribution of RT values (green) is compared to all non-centromeric regions of the genome (gray). Apart from transition zones (“CT”), which include ∼5Mb of the p- and q-arms flanking each centromeric region, all satellite families are biased toward late replication timing. However, the α-satellite higher-order repeats (“HOR”) are earlier-replicating than the large heterochromatic arrays (HSat2 and HSat3). RT values are for the lymphoblastoid cell line GM12878.

### Replication timing bias of repetitive sequence elements

Between the acrocentric p-arms and the centromeric regions, T2T-CHM13 adds 395Mb of densely annotated repeat-rich sequence whose replication timing has not been analyzed. Many of the annotated satellite sequences are relatively short (median: 7.25Kb) and neighbored by sequences of other satellite families (Figure 4a). Thus, we were interested to know whether these satellite families differed from one another in their replication timing: persistent patterns in replication timing of a family across multiple chromosome contexts could reflect some underlying property that controls when it replicates.

Indeed, satellite families did differ in both the median and range of replication timing values observed (Figure 4b). Replication timing values for non-repetitive sequence in these regions (annotated as “CT”) ranged from very early to very late, with a median somewhat later than the genome average (RT = -0.25 vs. -0.03). In contrast, each of the satellite sequence families was biased toward late replication – although none were exclusively late replicating. Notably, α-satellite HORs replicated earlier on average than HSat2 and HSat3, but later than HSat1. This is consistent with the notion that the active centromere is earlier replicating than its surrounding context, potentially to facilitate kinetochore loading onto both sister chromatids, at the appropriate time during S-phase. Furthermore, late replication of HSat2 and HSat3, evolutionarily related satellites that form large blocks of constitutive heterochromatin, suggests that they may comprise the later waves of replication observed by microscopy^2^.

### Replication dynamics within centromeric regions

Identifying the locations of replication timing peaks within centromeric regions allowed us to next ask about replication dynamics within these regions. We used two metrics to assess replication dynamics: the distance between consecutive replication timing peaks as a proxy for inter-origin distance, and the slope between replication timing peaks and valleys as a proxy for replication fork speed. We observed that inter-origin distances were slightly longer in centromeric regions relative to the rest of the genome (Figure 5a) and replication-timing slopes were slightly shallower (Figure 5b). While looking specifically within α-satellite HORs, these trends were more pronounced (Figure 5c,d). This could suggest that the active centromere poses a barrier to replication initiation and/or elongation, resulting in fewer origins firing and/or slower replication progression through these satellite arrays. However, there was substantial overlap between the distributions in all comparisons, indicating that many individual origins have similar dynamics in centromeric and non-centromeric regions. Thus, we favor the explanation that these differences are an artifact of the relatively sparser sequencing coverage of centromeric regions, resulting in an undercounting of centromeric peaks.

**Figure 5.**
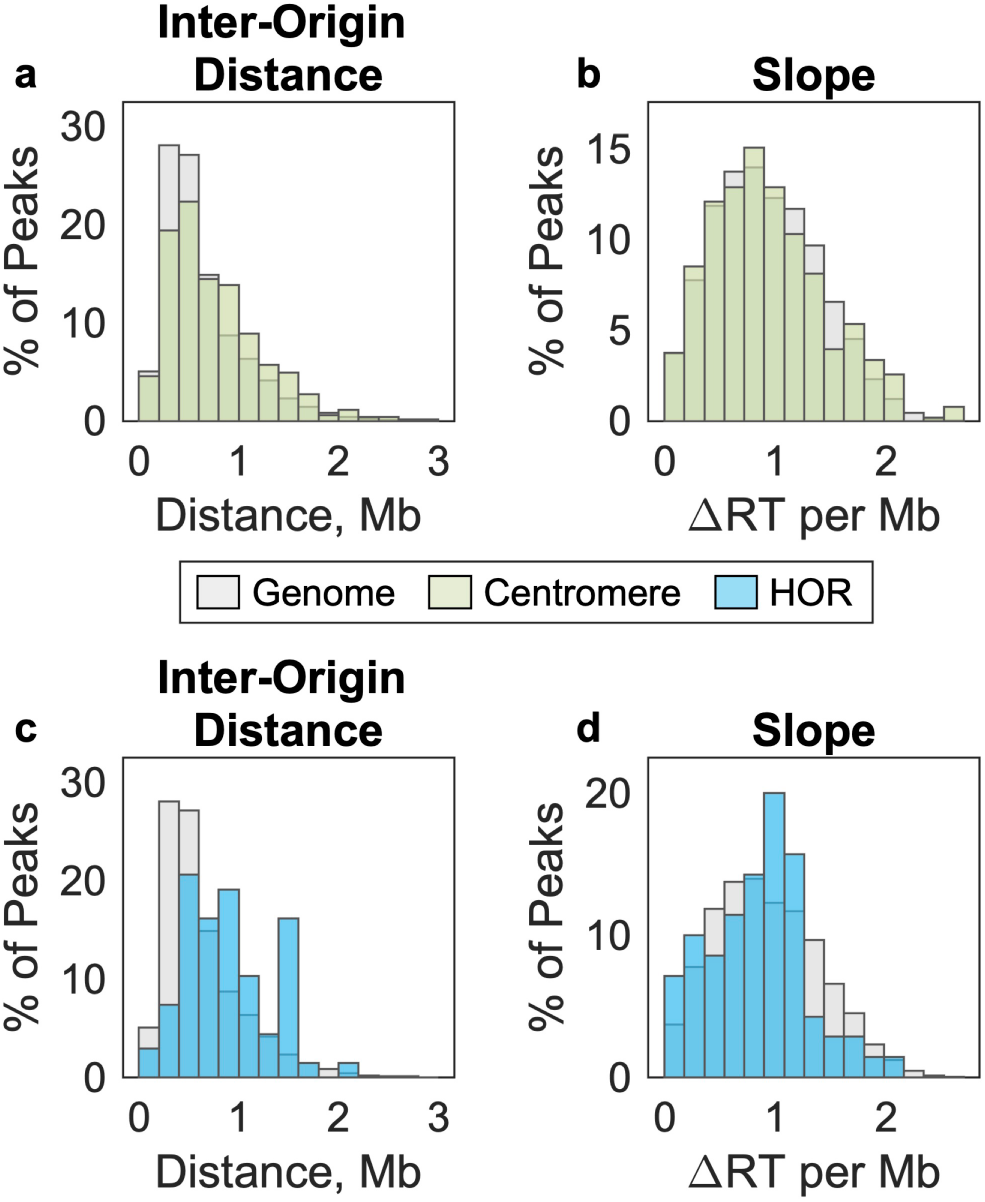
Replication timing (RT) peaks are not substantially different in centromeric regions than in the rest of the genome. **a, c** The distance between RT peaks was used as a metric of inter-origin distance. Inter-origin distances were slightly larger in centromeric regions (green, **a**) and α-satellite higher-origin repeats (blue, **c**), relative to the rest of the genome (gray). **b, d** RT profile slope was used as a proxy for replication fork speed. For each peak, the ascending and descending slopes are averaged. RT slopes were slightly shallower in centromeric regions (green, **b**) and α-satellite higher-origin repeats (blue, **d**), relative to the rest of the genome (gray). RT values are for the lymphoblastoid cell line GM12878.

### Centromeric replication timing varies consistently among cell lines

Finally, we considered differences between the five cell lines analyzed. Replication timing biases of individual satellite repeat families were consistent across cell lines (Figure 6a). Likewise, inter-origin distances (Figure 6b) and replication timing slopes (Figure 6c) were comparable. We had previously observed that there were differences in average centromeric replication timing between these cell lines, such that the average centromeric region in A2780 and HEK293T was early replicating and the average centromeric region in HCC1954 and HCC143 was late replicating^17^. Even though the replication timing profiles in these regions could not be “lifted over” between hg38 and T2T-CHM13, this trend was again observed in the T2T-CHM13 profiles (Figure 6d). Using T2T-CHM13, we were further able to analyze replication timing of individual centromeric regions in each cell line. We found that the trend observed on average reflected a persistent pattern across chromosomes within each cell line, rather than being driven by the replication timing of the larger centromeres (Figure 6e).

**Figure 6.**
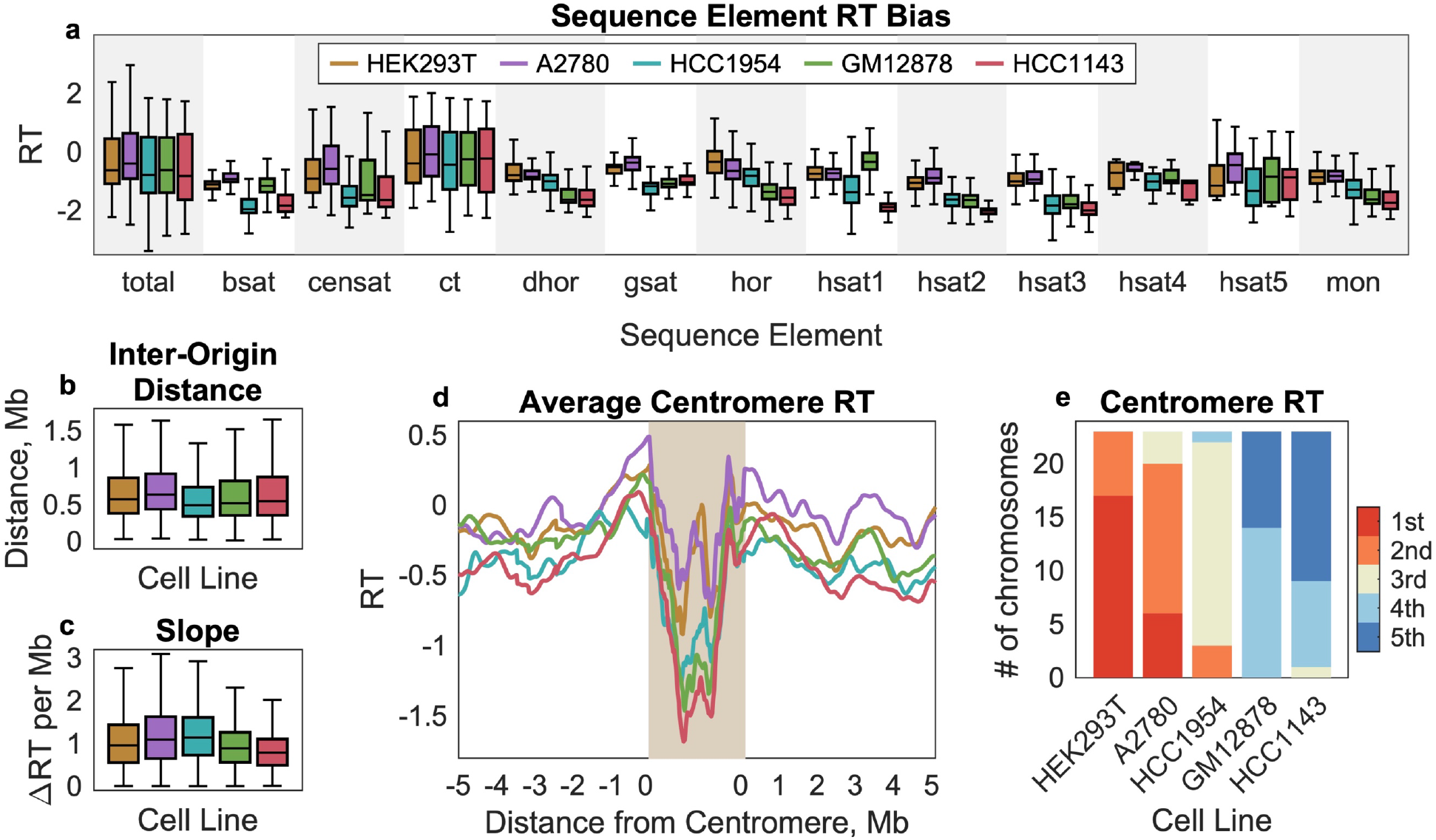
Variability in centromeric regions among cell lines persists across sequence elements and chromosomes. **a** The replication-timing bias for each centromeric sequence element type is compared across five cell lines. HEK293T and A2780, which have, on average, the earliest centromeric replication timing, are earlier replicating across many different sequence elements. Compare to Figure 4. **b, c** Inter-origin distance and RT slope are similar across cell lines. Compare to Figure 5. **d** Average replication-timing within centromeric regions and the flanking 5Mb on either side. For each chromosome, the centromeric region was divided into 100 equally spaced bins. HEK293T and A2780 have the earliest average centromeric replication, while GM12878 and HCC1143 have the latest. **e** Differences in centromere replication timing among cell lines are consistent across chromosomes. Each bar represents the number of times that a given cell line is the earliest, 2^nd^ earliest, 3^rd^ earliest, etc. HEK293T and A2780 are consistently the earliest replicating, while GM12878 and HCC1143 are consistently the latest replicating, and HCC1954 is consistently in between.

Taken together, our results indicate that the T2T-CHM13 genome assembly provides a reliable tool for inference of nearly gapless telomere-to-telomere human replication timing profiles. These newly profiled regions confirm that heterochromatin is typically (but not exclusively) late replicating and reveal differences in replication timing biases of satellite repeat families. Linear centromeric reference sequences enabled us to further confirm our prior findings that centromeres replicate in mid-to-late S phase, are not unusually late replicating relative to the rest of the genome, and that their timing of replication differs between cell lines. One biological mechanism that could potentially shape differences between cell lines is differential recruitment of the centromere-specific histone H3 variant CENP-A. Variation in HOR array length and sequence divergence has been shown to influence the competency of centromeric regions to recruit CENP-A^22^, and *in vitro* experiments suggest that depletion of CENP-A during S-phase results in replication fork stalling specifically at centromeres^23^. Thus, sequence and copy-number variation at centromeric regions among cell lines may alter the replication timing of individual chromosomes. However, by comparing centromeric regions within the same cell line, we demonstrate that earlier centromeric replication timing appears to be a global phenomenon impacting all chromosomes. An intriguing possibility is that centromeric replication is coordinated across chromosomes, perhaps by their nuclear localization: centromeres are strongly enriched for intrachromosomal interactions in budding yeast^24^ and centromere location within the nucleus has been implicated in the maintenance of pluripotency in human embryonic stem cell lines^25^. In that scenario, advancing the replication timing of one centromere could have the impact of altering global centromeric replication timing. To our knowledge, such a mechanism has yet to be described. Likewise, the consequences of divergent centromeric replication timing between cell lines remain unclear.

## Methods

### Preparation of whole genome sequence data

All sequence data analyzed in this study were previously published in Massey *et al*.^17^. Tissue culture, fluorescence-activated cell sorting, library preparation, and sequencing are detailed in that publication.

Sequencing reads were re-aligned to the human genome assembly T2T-CHM13 v1.1 with the Burrows-Wheeler Aligner maximal exact matches (BWA-MEM) algorithm (bwa v0.7.13). Sequence annotations are from Altemose *et al*.^20^ and were downloaded from the UCSC Genome Browser (University of California, Santa Cruz; “cenSatAnnotation” track). For acrocentric chromosomes, the p-arm boundary of the centromere was defined as 5Mb from the p-most HOR element. For chromosomes 1, 9, and 16, the q-arm boundary of the centromere was defined as 5Mb from the q-most HOR element.

### Replication timing profiles

Replication timing profiles were inferred by the S/G_1_ method described in Koren *et al*. (2012)^18^. Briefly, variable-size genomic bins were defined such that each bin had uniform coverage (200 reads) in the G_1_-phase library for a given cell line. Per-bin coverage was calculated for the corresponding S-phase library. The resulting profile was smoothed using a cubic smoothing spline (MATLAB function csaps, smoothing parameter 1 × 10^−16^), and normalized to an autosomal mean of 0 and standard deviation of 1.

## Acknowledgements

This work was supported by the National Institutes of Health (DP2-GM123495 to A.K.) and the National Science Foundation (MCB-1921341 to A.K.).

## Author Contributions

D.J.M. and A.K. conceptualized the project. D.J.M. performed analyses. D.J.M. and A.K. wrote the manuscript.

