## Supplementary Figures 1-3 for "Telomere-to-telomere human DNA replication timing profiles"

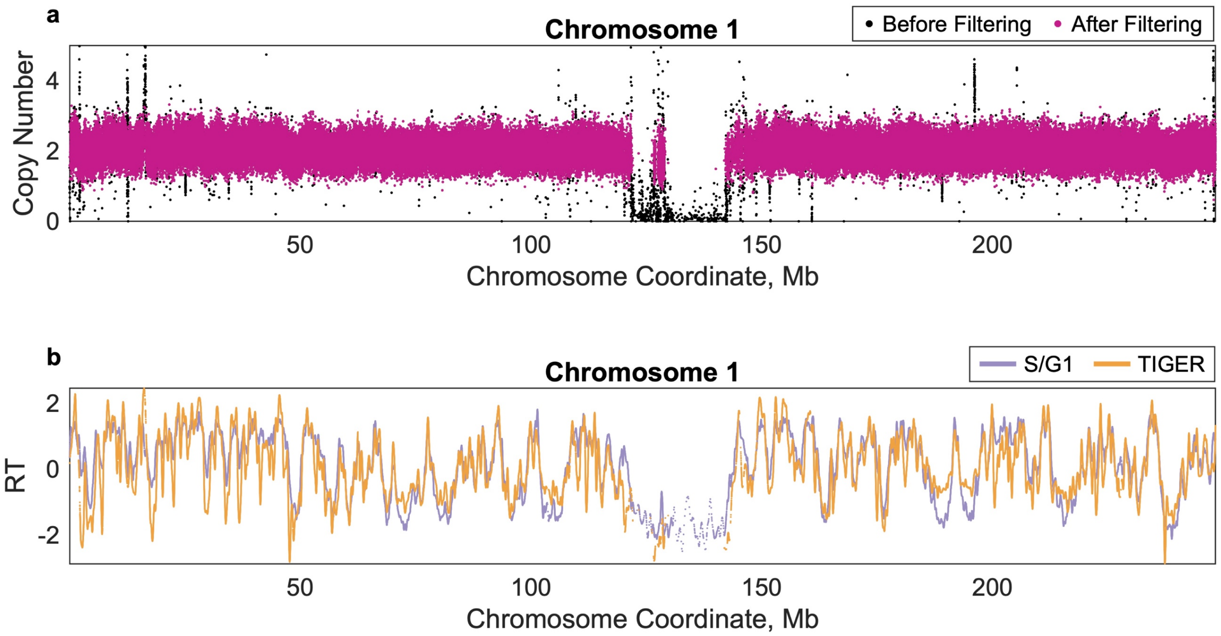


Supplementary Figure 1. **Replication timing analysis of highly repetitive regions requires a $G_1$-phase control sample. a** Mappability and GC-content corrected read counts for an asynchronous population of GM12878 cells before (black) and after (pink) filtering regions with abnormal copy-number estimates. The low coverage in the centromeric region is inadequately corrected even after accounting for these sequencing biases using TIGER[1]. **b** Similar replication timing profiles are obtained in non-repetitive regions of the genome between the S/$G_1$ and TIGER methods.

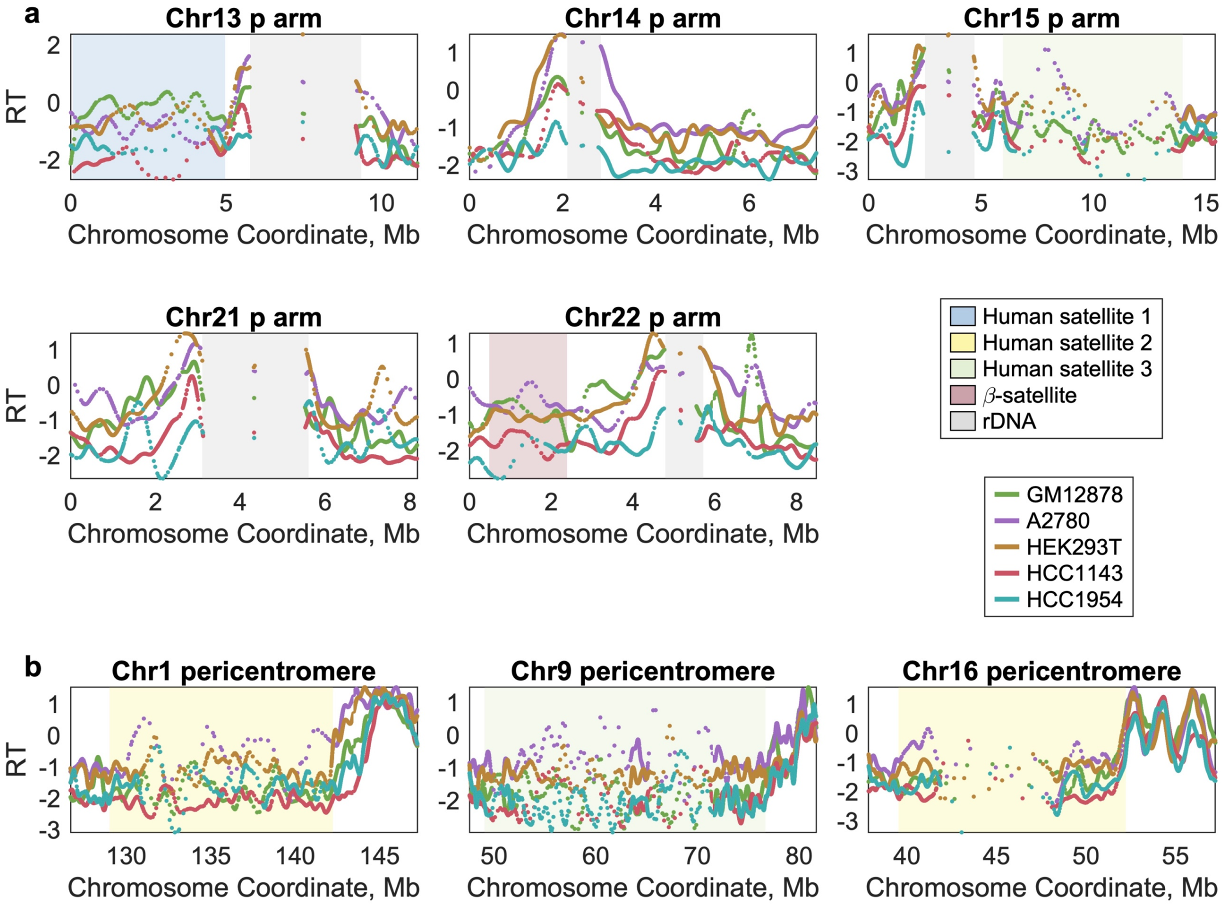


Supplementary Figure 2. **Replication timing (RT) of previously unresolved regions of the human genome for five cell lines.** Compare to Figure 2a,b.

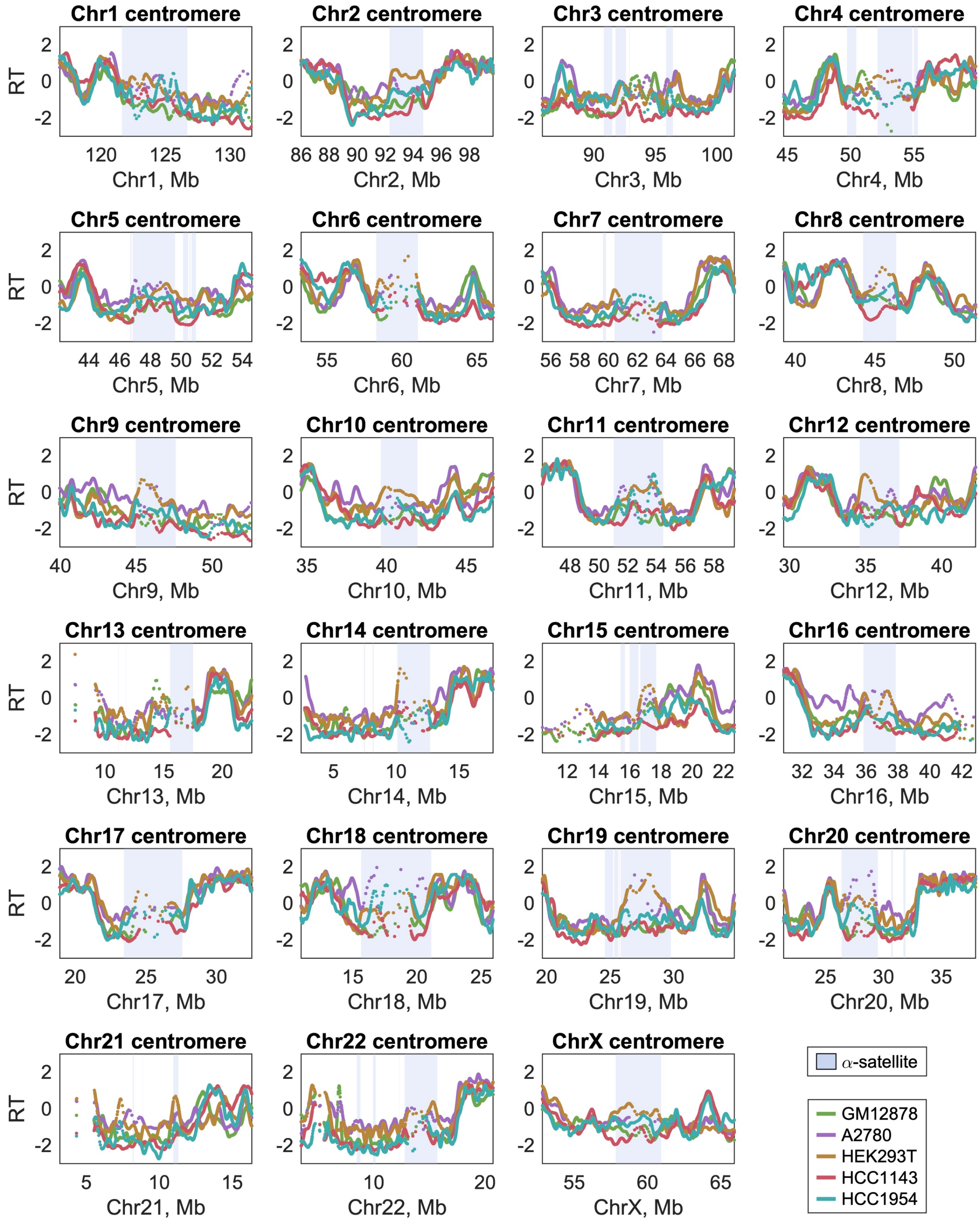


Supplementary Figure 3.**Centromere replication timing (RT) of all human autosomes and chromosome X for five cell lines.** Compare to Figure 3.
